# Disrupted Regional Dynamics of Structure-Function Connectivity Coupling in Euthymic Bipolar Disorder

**DOI:** 10.1101/2025.09.29.679408

**Authors:** Yi Ye, Kaitong Ma, Erni Ji, Xiaofen Zong, Maolin Hu, Zhen Liang, Gan Huang, Bharat B. Biswal, Xujun Duan, Li Zhang

## Abstract

**Background:** Bipolar disorder (BD) is characterized by persistent disturbances in emotional regulation and cognitive function, even during euthymia, yet its neural mechanisms remain unclear. Given evidence of structural and dynamic functional connectivity (dFC) abnormalities in BD, investigating dynamic structure-function connectivity coupling (SC-FC coupling) may provide insights into the pathophysiology of the disorder.

**Methods:** Diffusion tensor imaging and resting-state fMRI data were collected from 61 euthymic BD-I patients and 54 healthy controls (HC). Structural connectivity was derived using probabilistic tractography, and dFC was estimated with a sliding-window approach. SC-FC coupling was quantified as the extent to which structural features predicted dFC using multivariate linear regression. Abnormal SC-FC coupling in BD was assessed at both the network and regional levels, relative to HC. Over time, SC-FC coupling was categorized into high- and low-coupling states, and dynamic state analyses of dwell time, occurrence rate, and transition probabilities were performed in the affected brain regions.

**Results:** Network-level analyses indicated that BD exhibits lower SC-FC coupling in subcortical regions compared with HC. Regional analyses revealed significantly reduced dynamic SC-FC coupling in two thalamic subregions (medial posterior and rostral temporal) and in the right primary somatosensory cortex in BD compared to HC. Dynamic analyses further demonstrated thalamic instability: BD showed shorter dwell times in high-coupling states, increased occurrence of low-coupling states, and more frequent high-to-low coupling transitions than HC. These abnormalities suggest that BD is characterized by disrupted structure-function integration and unstable thalamocortical dynamics.

**Conclusion:** Our findings reveal regional abnormalities in SC-FC coupling dynamics in BD, indicating persistent dysfunction of structure-function integration during euthymia. Dynamic coupling measures may provide preliminary insights into potential biomarkers for disease monitoring.

## 1. Introduction

Bipolar disorder (BD) is a severe and chronic psychiatric illness characterized by recurrent episodes of mania, hypomania, and depression, leading to profound disturbances in mood, cognition, and daily functioning. Affecting more than 1% of the global population, BD is a leading cause of disability among young adults and imposes a substantial socioeconomic burden worldwide (Grande et al., 2016). Beyond its episodic nature, BD is associated with persistent cognitive deficits and functional impairments even during clinically stable periods. Advances in neuroimaging techniques, particularly functional magnetic resonance imaging (fMRI) and diffusion tensor imaging (DTI), have enabled the *in vivo* mapping of large-scale brain networks, revealing structural connectivity (SC) and functional connectivity (FC) patterns that are essential for understanding the neural mechanism of BD (Yen et al., 2023). Characterizing these network-level alterations holds promise for identifying biomarkers that may facilitate earlier diagnosis, refine disease subtyping, and guide targeted interventions.

Accumulating neuroimaging evidence indicates that BD involves widespread abnormalities in both functional and structural brain networks. Functional imaging studies consistently report aberrant activation and connectivity within key regions implicated in emotion processing, regulation, and reward, such as the thalamus, amygdala, orbitofrontal cortex, and anterior cingulate cortex (Zeng et al., 2023; Phillips et al., 2014; Wu et al., 2024). Resting-state FC analyses further reveal increased connectivity within emotion regulation-related networks, including the limbic network (LN), default mode network (DMN), and frontoparietal network, suggesting hypercoupling or imbalanced coordination between large-scale systems (Hu et al., 2023). Among the brain regions involved in emotion processing, the thalamus is particularly noteworthy. As a central relay station and integrative hub, the thalamus coordinates the flow of information between cortical and subcortical areas and plays a crucial role in emotional and cognitive regulation (Sherman et al., 2016; Hwang et al., 2017). Functional studies have revealed weakened connectivity within the striatum-thalamus circuit and altered coupling between the thalamus and limbic structures, changes that may underlie affective and cognitive disturbances (Teng et al., 2014). On the other hand, structural imaging studies of BD similarly demonstrate diffuse white matter microstructural abnormalities, including reduced fractional anisotropy and decreased integrity in prefrontal regions, potentially reflecting diminished neural signal conduction efficiency (Versace et al., 2008; Favre et al., 2019). Meanwhile, consistent evidence indicates that both structural and functional abnormalities are present in the thalamus in BD. Diffusion imaging studies have reported reduced fractional anisotropy in the anterior thalamic radiation, indicating compromised white matter integrity (Lin et al., 2011). These findings suggest that abnormalities in SC and FC may both contribute to the pathophysiology of BD, making it crucial to investigate their interrelationship.

The relationship between structural and functional connectivity, known as structure-function connectivity coupling (SC-FC coupling), offers a unified framework for linking the brain’s anatomical architecture to its dynamic functional organization (Honey et al., 2009; Fotiadis et al., 2024). SC-FC coupling is typically quantified by correlating SC matrices derived from diffusion imaging with FC matrices obtained from fMRI. This approach has been increasingly applied to psychiatric disorders, revealing altered SC-FC coupling patterns in BD and major depressive disorder, including enhanced structural “ rich-club” connectivity but reduced global coupling strength in BD (Zhang et al., 2019). However, most existing studies have relied on static coupling measures, overlooking the inherently dynamic nature of functional connectivity. Emerging evidence indicates that BD is characterized by pronounced temporal fluctuations and instability in functional networks, particularly reduced variability in dynamic functional connectivity (dFC) between cognitive and perceptual systems (Wang et al., 2020; Liu et al., 2021; Du et al., 2021). These observations suggest that static measures may fail to capture the changes in brain networks across different states. Given that the brain is a system that is constantly changing, it is therefore important to investigate dynamic SC-FC coupling, which explicitly incorporates the intrinsic temporal dynamics of the brain. This approach provides a more comprehensive reflection of the spatiotemporal characteristics of brain networks and allows for the detection of state-dependent changes that static measures may ignore.

Based on these observations and evidences, it is reasonable to hypothesize that BD patients exhibit abnormal SC-FC coupling in certain key brain regions, such as the thalamus, which may reflect focal network disruptions contributing to the broader instability of brain connectivity observed in the disorder. In order to test our hypothesis, we propose a novel framework aimed at comprehensively characterizing the dynamic properties of SC-FC coupling in BD from multiple perspectives. By combining resting-state fMRI (rs-fMRI) and DTI, we constructed dFC matrices using a sliding-window approach and obtained SC matrices through probabilistic tractography. Dynamic SC-FC coupling was quantified at both the network and regional levels using multivariate linear regression, and temporal stability was assessed through autocorrelation analysis. Based on the regional-level results, we identified key brain regions exhibiting significant coupling abnormalities between BD patients and healthy controls. Further dynamic state analyses, including measures of dwell time, occurrence rate, and transition probabilities, were performed to explore the time-varying coupling patterns in these regions. Our findings reveal significant abnormalities in SC-FC coupling across multiple key brain regions in BD patients, particularly in dynamic coupling patterns, showing temporal instability and changes in spatial distribution. These abnormalities provide new insights into the dynamic interplay between brain structure and function in psychiatric disorders.

## 2. Materials and Methods

### 2.1 Participants

Fig. 1 presents the flow chart of the analysis conducted in this study. The study enrolled 61 euthymic, medicated BD-I patients (30 males, 31 females; mean age ± SD = 31.85 ± 7.97 years) and 54 healthy controls (HC; 26 males, 28 females; mean age ± SD = 29.78 ± 7.17 years). BD patients were recruited from the Shenzhen Mental Health Center, and HCs through public advertisements. All procedures performed in this study were approved by the Human Research Ethics Committee of the Shenzhen Mental Health Center (2015-K015-03), and the trial was registered in the Chinese Clinical Trials Registry (MR-11-24-048384). All procedures were conducted in accordance with the ethical standards outlined in the Declaration of Helsinki. All participants provided written informed consent. BD diagnoses were confirmed using the Structured Clinical Interview for DSM-IV (SCID) (Gorgens, 2011), with all patients meeting the criteria for BD-I, which requires at least one manic episode, possibly accompanied by depressive episodes. Patients were on stable lithium and valproate treatment for at least three months and were euthymic at the time of scanning, as confirmed by the Hamilton Depression Rating Scale (Hamilton, 1967) (Chinese version) and the Young Mania Rating Scale (Young et al., 1978) scores (Chinese version). Inclusion criteria for BD required MRI eligibility, no current mood episodes, and no hospitalization in the past six months. HCs had no psychiatric or neurological history, no significant head trauma, no first-degree family history of major psychiatric disorders, dementia, or intellectual disability, and no recent substance dependence. A non-clinical SCID interview (First et al., 2004) confirmed the absence of psychiatric or neurological conditions in HCs.

**Fig. 1.**
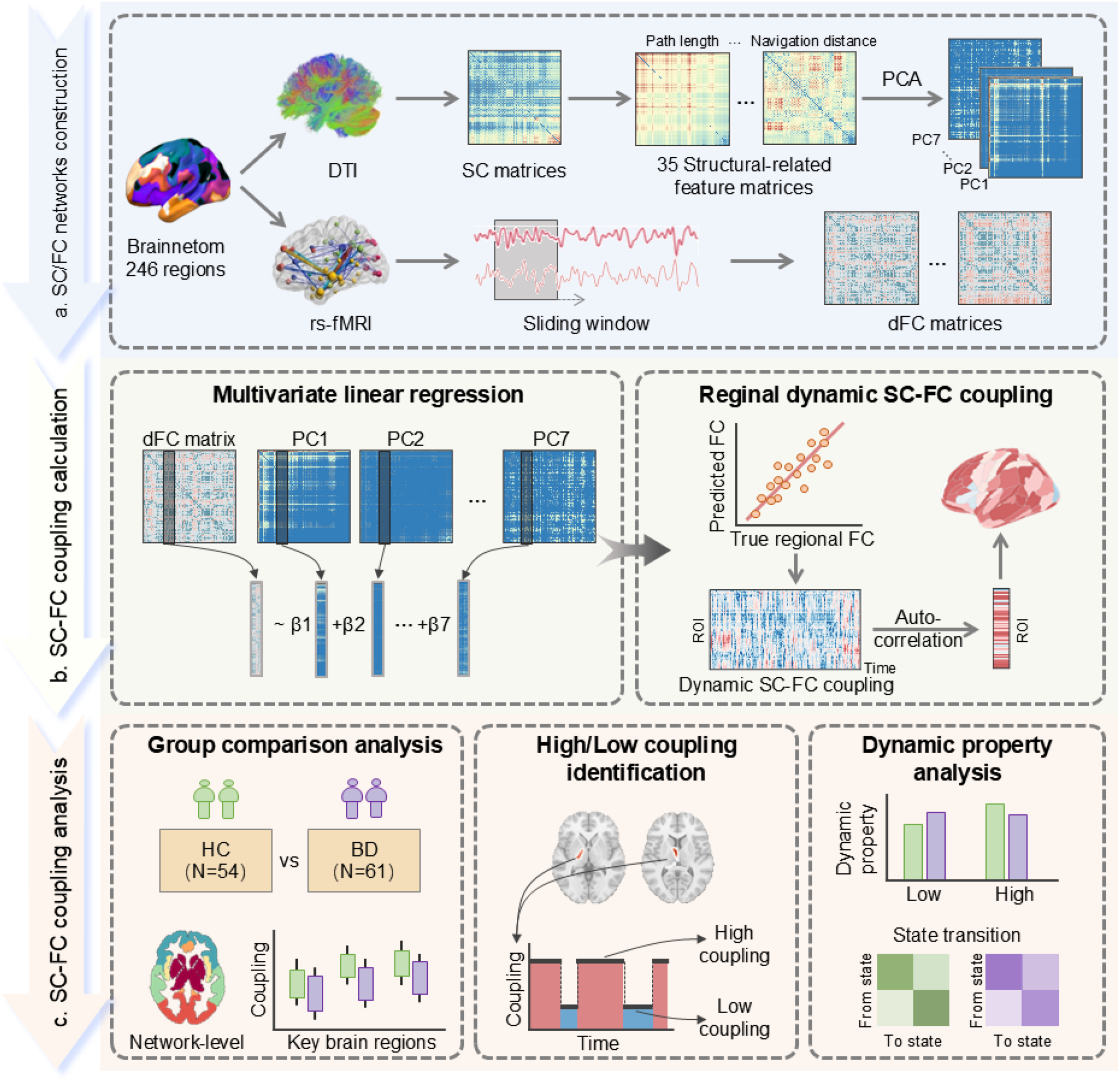
Workflow of SC-FC coupling analysis. (a) SC/FC network construction: DTI and rs-fMRI data were parcellated into 246 ROIs using the Brainnetome atlas. SC matrices were generated from DTI-based tractography, and 35 structure-related feature matrices were further derived. PCA was applied to extract the first *k* components explaining >80% variance. dFC matrices were obtained from rs-fMRI data using a sliding-window approach. (b) SC-FC coupling calculation: For each ROI and time window, regional FC values were regressed against the retained structural components in a multivariate linear regression model. SC-FC coupling was defined as the Pearson correlation between predicted and observed FC values, while autocorrelation coefficients quantified the temporal stability of ROI-level coupling dynamics. (c) SC-FC coupling analysis: ROI-level ANCOVA was performed between BD and HC groups to identify key abnormal regions with disrupted coupling. Coupling patterns of these regions were further classified into high and low states based on global mean values, and dynamic properties, including state transition counts, mean dwell time, occurrence rate, and transition probabilities, were calculated. Group differences in these dynamic characteristics were examined using two-way ANCOVA, with diagnostic group and coupling state as fixed factors.

### 2.2 Data Preprocessing

After clinical assessment, all participants underwent imaging on a Siemens 3T Trio MRI scanner equipped with a 12-channel head coil to acquire rs-fMRI and DTI data. Resting-state fMRI data were obtained using a standard gradient-echo echo-planar imaging (EPI) sequence with the following parameters: 31 oblique slices, repetition time (TR) = 2000 ms, echo time (TE) = 30 ms, field of view (FOV) = 240 × 240 mm^2^, flip angle = 90°, matrix size = 64 × 64, voxel size = 3 × 3 × 5 mm^3^, and a total of 246 volumes. During the entire scan, participants were instructed to keep their eyes open and remain awake. Preprocessing of the rs-fMRI data was conducted in MATLAB R2021a using the DPABI toolbox (http://rfmri.org/dpabi/) (Yan et al., 2016). For each subject, the first 10 time points were discarded to minimize the effects of magnetic field instability. The remaining images were then subjected to slice-timing correction and motion correction. To further reduce non-neural noise, time points with head motion exceeding 0.2 mm were excluded and regressed out as separate covariates. In addition, white matter and cerebrospinal fluid signals were regressed out as nuisance covariates. Subsequently, the functional images were normalized to the Montreal Neurological Institute (MNI) standard space, resampled to 3 × 3 × 3 mm^3^ voxels, spatially smoothed with a 4 mm full-width at half-maximum (FWHM) Gaussian kernel, and band-pass filtered (0.01-0.1 Hz) to improve the signal-to-noise ratio.

DTI preprocessing was conducted in FSL following the procedures described in (Kim et al., 2022). Eddy current and head motion corrections were first applied to the raw DTI images, after which the Brain Extraction Tool (BET) (Smith, 2002) was used to remove non-brain tissue and generate a brain mask. Diffusion parameters were then estimated for each voxel using BedpostX (Behrens et al., 2007) to model fiber crossing within voxels. Finally, the diffusion-weighted images were normalized to the MNI space to ensure spatial alignment across subjects.

### 2.3 SC and FC Networks Construction

After preprocessing of the rs-fMRI and DTI data, the Brainnetome Atlas (Fan et al., 2016) was applied to parcellate the whole brain into 246 regions of interest (ROIs), each corresponding to a specific anatomical area. For rs-fMRI data, the blood-oxygen-level-dependent (BOLD) time series of each ROI was extracted. For DTI data, the number of streamlines between each pair of ROIs was estimated using probabilistic tractography.

Functional connectivity analysis was conducted using a sliding-window approach. A rectangular window of 50 TRs with a step size of 1 TR was applied to the BOLD time series, producing overlapping temporal segments. Within each window, Pearson correlation coefficients were computed between the time courses of all ROI pairs, followed by Fisher’s Z-transformation to improve normality. This procedure yielded, for each subject, a dFC matrix of size 246 × 246 × 236, where 246 is the number of ROIs and 236 is the number of temporal windows.

Structural connectivity analysis was performed using FSL. Diffusion-weighted images were processed with BedpostX to estimate the probability distribution of fiber orientations, and co-registered with T1-weighted images to the standard MNI space using linear and nonlinear transformations. Probabilistic tractography (ProbtrackX) was then applied to calculate the number of streamlines between each pair of ROIs, generating the SC matrix. Streamline counts were normalized by the average volume of the corresponding seed and target ROIs, and connections with a coefficient of variation below the 25th percentile were excluded. The matrix was symmetrized by averaging its upper and lower triangular components, resulting in a sparse, undirected, symmetric SC matrix. In parallel, Euclidean distances between ROIs were computed from centroid coordinates in the brain atlas. Based on the SC matrices, 34 structure-related feature matrices were derived using a communication model similar to that described by (Popp et al., 2024), including matching index, cosine similarity, path length, search information, communicability, mean first passage time, navigation, flow graph, and path transitivity measures. Interested reader may refer to the supplementary documents for more details about these 34 feature matrices.

### 2.4 SC-FC Coupling Calculation

SC-FC coupling calculation was carried out by performing principal component analysis (PCA) on 36 feature matrices, including 34 aforementioned structure-related feature matrices, the SC matrix and the Euclidean distance matrix. The first *k* principal components explaining over 80% of the variance were retained. For each subject and each time window, a multivariate linear regression model was constructed, with independent variables comprising the FC connection strengths of a given ROI (one column of the dFC matrix) and the *k* SC principal components for the same ROI. The dependent variable was the predicted FC value for that ROI. Actual FC values were z-score normalized by column to control for scale differences across ROIs. The Pearson correlation coefficient between predicted and actual FC values was defined as the SC-FC coupling value for that ROI. To further characterize the temporal properties of coupling, autocorrelation coefficients with a lag of 15 were computed from the coupling time series to quantify the stability and delayed dependencies of dynamic SC-FC coupling. The lag was chosen based on prior dynamic functional connectivity studies (Hutchison et al., 2013; Preti et al., 2017) to ensure a balance between sensitivity and robustness.

### 2.5 SC-FC Coupling Analysis

We first investigate large-scale network-level abnormalities. All ROIs defined by the Brainnetome atlas were grouped into seven canonical networks: frontal, temporal, parietal, insular, limbic, occipital, and subcortical. For each subject, the SC-FC coupling autocorrelation coefficients of ROIs belonging to the same network were aggregated to compute a network-level mean coupling value, reflecting the overall strength of structure-function alignment within the network. These measures were derived separately for each of the seven networks. Subsequently, analysis of covariance (ANCOVA) was performed to compare the bipolar disorder (BD) group with the healthy control (HC) group. Years of education were included as a covariate to control for potential confounding effects, given the significant difference between the two groups. Statistical significance was set at *p* < 0.05, with false discovery rate (FDR) method for multiple comparisons correction.

Next, we examined the temporal properties of SC-FC coupling at the ROI level. Autocorrelation coefficients for each ROI were subjected to one-way ANCOVA with years of education as a covariate, and group comparisons between BD and HC were corrected for multiple comparisons using the FDR method. ROIs with FDR-corrected *p* < 0.05 were defined as key regions with abnormal dynamic coupling. These regions indicate focal disruptions in the temporal correspondence between structural and functional connectivity, reflecting potential network vulnerabilities within the broader connectome. They were selected as primary targets for subsequent analyses of coupling instability, allowing us to characterize how abnormalities evolve over time in the most affected areas. For visualization, Cohen’s d effect sizes of significant ROIs were calculated and mapped onto the Brainnetome template to illustrate cortical and subcortical distributions of coupling alterations.

To characterize the dynamic coupling properties of the key regions with abnormal dynamic coupling, we extracted the coupling value matrix of identified abnormal regions that exhibited significant between-group differences for each subject. The global mean coupling value across all participants and all the time windows was then calculated and used as a threshold to define coupling states: values above the mean were classified as the high-coupling state, and values below the mean were classified as the low-coupling state. For each subject, dynamic properties of these coupling states were computed, including the number of state transitions, mean dwell time, state occurrence rate, and the state transition probability matrix. A two-way ANCOVA was conducted, with diagnostic group (BD vs. HC) and coupling state as fixed factors and years of education as a covariate, to examine the main and interaction effects on these temporal characteristics.

## 3. Results

### 3.1 Demographic Characteristics

The final sample consisted of 61 participants with BD and 54 HC. Demographic characteristics are summarized in Table 1. No significant between-group differences were observed in age or sex distribution, indicating that the groups were well matched on these variables. However, years of education differed significantly between BD and HC participants, which could potentially influence cognitive and neuroimaging measures. To minimize the impact of this disparity and improve the validity of group comparisons, years of education were included as a covariate in all subsequent statistical analyses.

**Table 1.** Demographic characteristics of all subjects.

| No.of subjects<br>(male/female) | BD |  | HC |  | P value |
| --- | --- | --- | --- | --- | --- |
|  | 61 (30/31) |  | 54 (26/28) |  | 1.000<sup>a</sup> |
|  | Mean<br>(SD) | Min-Max | Mean<br>(SD) | Min-Max | BD vs. HC |
| Age (years) | 31.85<br>(7.91) | 18-54 | 29.78<br>(7.10) | 22-47 | 0.147<sup>b</sup> |
| Education (years) | 13.33<br>(2.73) | 8-18 | 14.65<br>(2.40) | 8-21 | 0.005<sup>b</sup> |
| HAMD Score | 1.87<br>(1.99) | 0-7 | -- | -- | -- |
| YMRS Score | 0.62<br>(1.12) | 0-6 | -- | -- | -- |

Note: HAMD: Hamilton Depression Scale; YMRS: Young Mania Rating Scale. BD: Patients with bipolar disorder. HC: Healthy controls. <sup>a</sup> Pearson Chi-square test. <sup>b</sup> Independent two-sample t-test.

### 3.2 Aberrant Dynamic SC-FC Coupling in Thalamic and Somatosensory Regions

PCA revealed that the first seven principal components accounted for more than 80% of the explainable variance and were retained for subsequent SC-FC coupling calculations. In the network-level analysis, among the seven canonical networks, no significant group differences were observed for the all seven networks. Though all results did not reach statistical significance after FDR correction, the subcortical network showed a trend toward significance group differences in mean of SC-FC coupling autocorrelation coefficients. Specifically, the BD group exhibited lower mean coupling compared with the HC group (*pFDR* = 0.0573). For the ROI-level analysis, three brain regions showed significant between-group differences after FDR correction, involving left medial pre-motor thalamus (mPMtha, Cohen’s d = 0.696; 95% CI: 0.007 to 0.010; *p*FDR = 0.0353), left left rostral temporal thalamus (rTtha, Cohen’s d = 0.752; 95% CI: −0.009 to 0.007; *p*FDR = 0.0319) and right primary somatosensory cortex (A1/2/3 upper limb, head and face region (ulhf), Cohen’s d = 0.696; 95% CI: −0.006 to 0.011; *p*FDR = 0.0353). These regions span both subcortical and cortical structures that are integral to somatosensory processing, motor control, and the relay of information between cortical and subcortical systems (Alitto et al., 2003).

Effect sizes (partial eta squared) for all brain regions were converted to Cohen’s d and mapped onto the Brainnetome atlas to visualize the cortical and subcortical distribution of group differences (Fig. 2a). Post hoc comparisons further confirmed that mean SC-FC coupling in all three regions was significantly reduced in the BD group compared with the HC group (Fig. 2b), indicating a weakened correspondence between structural and functional connectivity in both thalamic and somatosensory relay circuits.

**Fig. 2.**
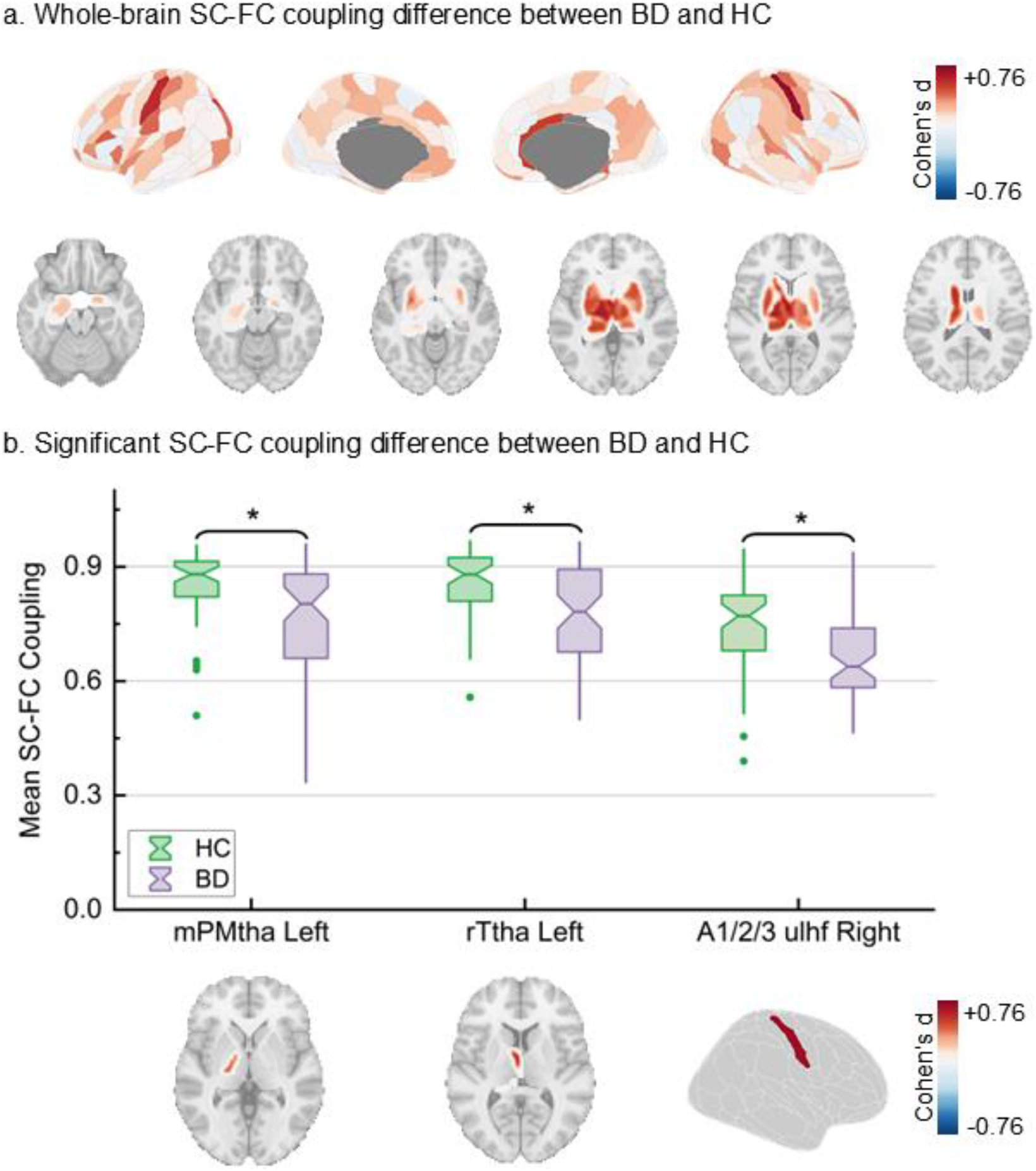
Group differences in SC-FC coupling between BD and HC. (a) ROI-level distribution of SC-FC coupling differences mapped using Cohen’s d. (b) Key regions showing significant reductions in BD compared with HC, including the left medial pre-motor thalamus (mPMtha), and left rostral temporal thalamus (rTtha) and right primary somatosensory cortex (A1/2/3 ulhf).

### 3.3 Instability of Thalamic SC-FC Coupling Dynamics

Based on the classification of the thalamic coupling time series, all time windows for each participant were categorized into high- and low-coupling states. The results of the two-way ANCOVA comparing state transition counts, state dwell time, and state occurrence rate are presented in Fig. 3. No significant group differences were observed in the number of state transitions (Fig. 3a; *p* > 0.05). However, the HC group demonstrated a significantly longer mean dwell time in the high-coupling state compared with the BD group (Fig. 3b; *p* = 0.035; Cohen’s d = 0.441). Significant between-group differences were also found in state occurrence rates (Fig. 3c). The BD group exhibited a higher occurrence rate of the low-coupling state (*p* = 0.004; Cohen’s d = −0.645) and a lower occurrence rate of the high-coupling state (*p* = 0.004; Cohen’s d = 0.645) relative to the HC group. Analysis of state transition probabilities (Fig. 3d) further revealed that the BD group had an increased likelihood of remaining in the low-coupling state (*p* = 0.050; Cohen’s d = −0.467) and a markedly greater probability of transitioning from the high-to the low-coupling state (*p* = 0.001; Cohen’s d = −0.665). In contrast, the HC group was more likely to maintain the high-coupling state (*p* = 0.001; Cohen’s d = 0.665).

**Fig. 3.**
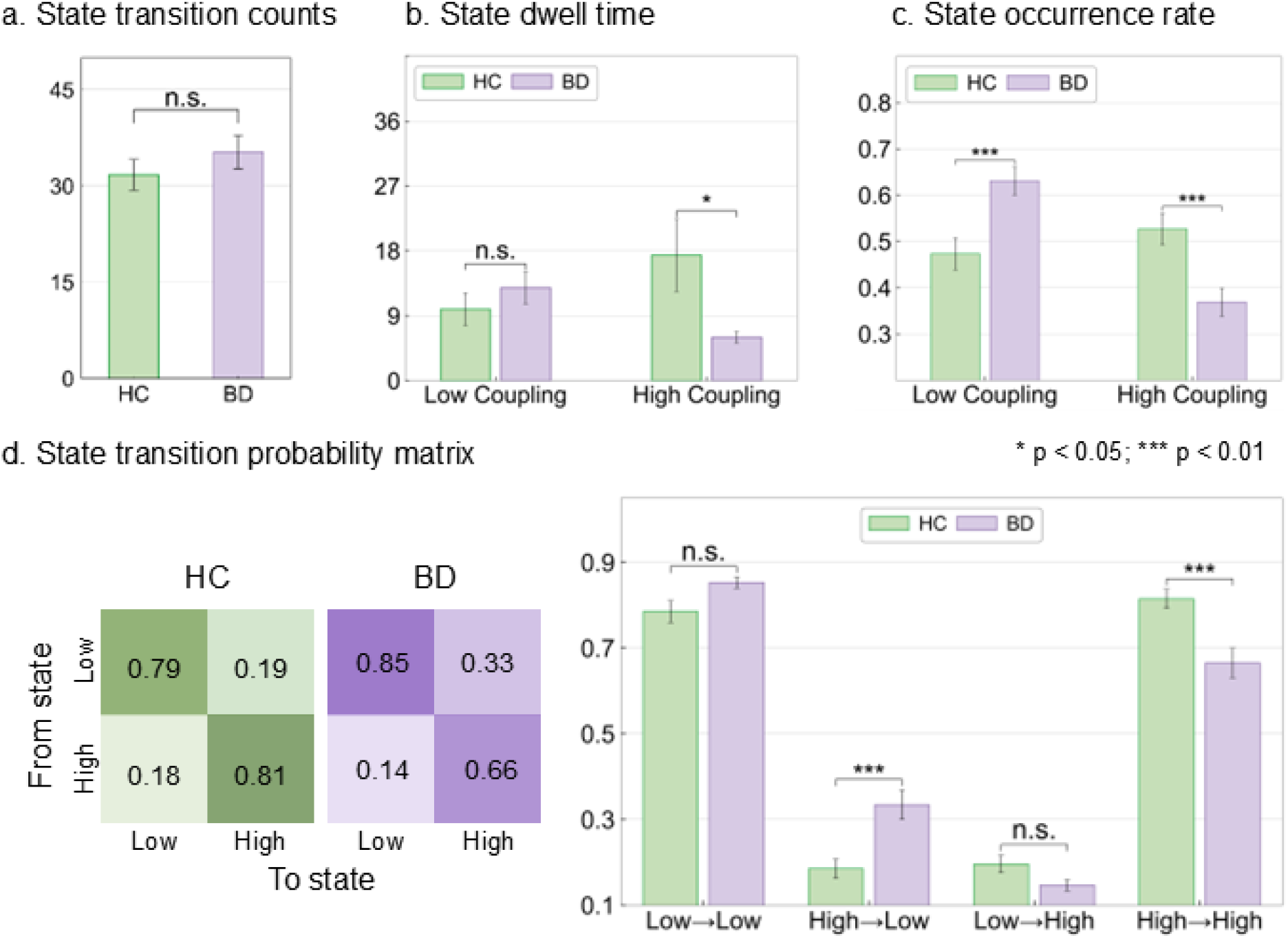
Dynamic properties analysis of thalamic SC-FC coupling. (a) No significant group difference was found in state transition counts between BD and HC. (b) BD patients showed significant shorter dwell time in high-coupling states. (c) BD patients exhibited higher occurrence of low-coupling states and lower occurrence of high-coupling states. (d) State transition probabilities indicated that BD patients were more likely to remain in or transition to low-coupling states, whereas HC tended to maintain high-coupling states.

## 4. Discussion

In this study, we combined rs-fMRI and DTI with a regional-level SC-FC coupling framework to investigate dynamic coupling alterations in euthymic BD-I patients. Using the Brainnetome atlas and a model-based approach to integrate structural connectivity features, we found that BD patients exhibited significantly reduced SC-FC coupling in three brain regions compared with HC: the left mPMtha, the left rTtha, and the right A1/2/3 ulhf in the primary somatosensory cortex. Based on these coupling abnormalities group comparison, given the critical role of the thalamus as a hub for emotional regulation, we focused on its dynamic coupling. The results showed that BD patients exhibited greater instability in high-coupling states, characterized by shorter dwell times, higher occurrence rates of low-coupling states, and an increased likelihood of transitioning from high-to low-coupling states. These findings indicate that both thalamic circuits and primary somatosensory cortex in BD show impaired structure-function integration and disrupted temporal stability, suggesting a multi-level network dysfunction even in the absence of acute mood symptoms.

### 4.1. Thalamic and Somatosensory SC-FC Coupling Alterations in BD

The SC-FC coupling analysis revealed significantly reduced coupling in multiple cortical and subcortical regions in BD compared with HC, even during the euthymic phase. Affected regions included two thalamic subregions associated with motor (mPMtha) and anterior temporal (rTtha) functions, as well as the right primary somatosensory cortex (A1/2/3 ulhf). These regions are critically involved in sensorimotor processing, emotional regulation, and the integration of cognitive and affective information, which are domains frequently impaired in BD patients (Kropf et al., 2019; Pantazatos et al., 2014; Teng et al., 2014). The thalamus acts as a central hub mediating information flow between subcortical structures and distributed cortical networks, including sensory, motor, and limbic systems (Greene et al., 2020; Zhuang et al., 2023). The primary somatosensory cortex plays a central role in integrating bodily sensations with higher-order cognitive and emotional processes; its reduced coupling may contribute to altered sensory perception and psychomotor regulation commonly observed in BD. Consistent with prior systematic reviews, BD has been associated with impaired regulation within subcortical striatal-thalamic-prefrontal networks and the associated limbic modulating regions, a deficit thought to underlie core emotional dysregulation in the disorder (Strakowski et al., 2005). The observed reduction in SC-FC coupling may reflect either compensatory structural reorganization in response to functional network imbalance or a diminished capacity for integration between structural pathways and functional activity (Lee et al., 2017). Together, these findings support the view that BD involves multi-level disruptions in cortical-subcortical communication, particularly within sensory-motor and thalamocortical circuits, which may serve as potential neurophysiological markers and persist beyond acute mood episodes.

Beyond the multi-level abnormalities, we conducted dynamic analyses focusing on the thalamus, given its central role in integrating sensory information, regulating cortico-subcortical communication, and supporting cognitive-emotional processing. The results revealed that BD patients exhibited significant instability in SC-FC coupling states. This instability was characterized by frequent transitions from high-coupling states, representing relatively stable and highly integrated network configurations, to low-coupling states that are unstable and decoupled. Compared with HC, BD patients showed a significantly higher occurrence rate of low-coupling states, a lower occurrence rate of high-coupling states, and shorter average dwell times in high-coupling states, along with a greater likelihood of transitioning from high to low coupling. The thalamus is a central relay structure regulating information flow between widespread cortical and subcortical networks, and its dynamic flexibility is essential for maintaining efficient integration across functional systems (Hwang et al., 2017). Prior studies have reported abnormal dynamic transition patterns in the functional connectivity networks of BD patients (Du et al., 2021), suggesting that such frequent state switching may reflect a heightened vulnerability of thalamocortical circuits to disengage from integrated network configurations. This instability may impair sensory gating and attentional control (Anticevic et al., 2014; Woodward et al., 2012), making BD patients more vulnerable to emotional imbalance and behavioral fluctuations when confronted with internal or external stimuli. On a deeper level, disrupted SC-FC coupling dynamics may be closely linked to the subtle yet persistent neurocognitive deficits in BD, ultimately contributing to emotional dysregulation and greater affective variability (Tiihonen et al., 2005; Renaud et al., 2012; Pomarol-Clotet et al., 2015). Collectively, these findings highlight that dynamic measures of structure-function coupling capture critical aspects of network dysfunction in BD that may not be detectable through static analyses alone.

### 4.2. Methodological Consideration

Previous studies have shown that functional connectivity disruptions are a key mechanism underlying cognitive and emotional processing deficits in patients with BD, with abnormalities in dynamic functional connectivity being closely associated with executive dysfunction (Nguyen et al., 2017). Therefore, this study introduces a dynamic perspective to compute SC-FC coupling by integrating dynamic functional and structural connectivity within a unified framework, and further evaluates the temporal stability of coupling states. Several methodological strengths enhance the robustness of our findings. First, our approach takes into account the inherent dynamic nature of brain activity by introducing a sliding-window method to characterize the time-varying features of structural-functional coupling. This enables us to investigate the dynamic properties of BD patients from a temporal perspective. Second, by integrating structural and functional imaging modalities within the SC-FC coupling framework, we were able to capture multi-level aspects of brain network organization that are not accessible through single-modality approaches. Third, our analysis incorporated both network-level and regional coupling measures, enabling us to detect not only persistent coupling abnormalities but also temporal instabilities in thalamocortical circuits.

Our findings align with and extend previous neuroimaging studies reporting disrupted thalamocortical connectivity in BD. Prior research using resting-state fMRI has consistently demonstrated altered thalamic functional connectivity patterns in both manic and euthymic phases (Öngür et al., 2010; Anticevic et al., 2014), while diffusion-based studies have identified microstructural abnormalities in thalamic radiations and fronto-striatal tracts (Favre et al., 2019; Yu et al., 2025). Although earlier work has examined dynamic functional connectivity alterations in BD (Du et al., 2021; Rashid et al., 2014), our results reveal that coupling dynamics between structural and functional networks provide complementary information to purely functional measures. Notably, the observed shift toward low-coupling states in BD may partially explain previous reports of reduced network integration and increased modular segregation in functional connectivity studies (Bai et al., 2020). Differences across studies may also be attributable to methodological factors, such as the choice of brain parcellation schemes, sliding window parameters, and the inclusion of patients with varying clinical characteristics and medication regimens. By using the Brainnetome atlas and controlling for education as a covariate, our approach reduces potential confounding and offers a more fine-grained characterization of coupling abnormalities.

From a clinical perspective, the identification of the instability of thalamic coupling states in BD offers potential avenues for biomarker development. Such coupling metrics may serve as objective neurophysiological indicators for detecting subtle network dysfunctions that persist during euthymia, potentially aiding in early diagnosis, monitoring disease progression, and predicting relapse risk (Hajek et al., 2013; Kambeitz et al., 2017). Moreover, dynamic coupling measures could provide a novel target for evaluating the effects of pharmacological and non-pharmacological interventions, such as mood stabilizers, cognitive remediation, or neuromodulation techniques. These measures can be used to assess whether treatment restores more stable and integrated network configurations. (Goya-Maldonado et al., 2016). Theoretically, our findings support a systems-level model of BD in which persistent disruptions in thalamocortical circuits, contribute to core deficits in emotional regulation, sensory processing, and cognitive control (Phillips et al., 2008; Strakowski et al., 2012). By demonstrating that regional dynamic coupling abnormalities, this study emphasize the importance of considering temporal network dynamics in addition to static connectivity when investigating the pathophysiology of mood disorders (Preti et al., 2017; Hutchison et al., 2013).

### 4.3. Limitations and Future Directions

Several limitations of this study should be noted. First, all patients included in this study were individuals with BD-I in the euthymic phase, and those in the acute phase were not considered. However, neural function and dynamic SC-FC coupling characteristics in acute-phase patients may differ substantially from those in the euthymic phase. Acute episodes are often accompanied by more pronounced fluctuations in brain network dynamics and abnormal connectivity patterns, which may reflect critical stages of disease progression (Martino et al., 2016). Therefore, future studies should include BD patients across different clinical states to systematically elucidate abnormalities in dynamic SC-FC coupling and their underlying mechanisms at various stages of the disorder. Medication effects, while minimized by recruiting euthymic and clinically stable patients, cannot be completely ruled out given the known influence of mood stabilizers on brain connectivity (Syan et al., 2018). In addition, our sample size, although comparable to prior neuroimaging studies in BD, may limit statistical power for detecting more subtle group differences. Future research should adopt longitudinal designs to track changes in SC-FC coupling across mood states and disease progression, examine the potential predictive value of dynamic coupling measures for relapse risk, and explore whether targeted interventions can restore coupling stability. Incorporating multimodal neuroimaging with cognitive, behavioral, and clinical assessments will further elucidate the links between network-level disruptions and functional outcomes in BD.

## 5. Conclusion

In conclusion, this study demonstrates that BD is characterized by significant instability in thalamic and reduction in primary somatosensory SC-FC coupling dynamics, even during euthymia. These abnormalities likely reflect persistent disruptions in structure-function integration that contribute to deficits in emotional regulation, sensory processing, and cognitive control. By combining network-level and regional dynamic coupling analyses within a multimodal neuroimaging framework, our findings provide novel insights into the multi-level pathophysiology of BD and identify potential neurophysiological markers for clinical monitoring.

## Ethical approval

All procedures performed in this study were approved by the Human Research Ethics Committee of the Shenzhen Mental Health Center (2015-K015-03), and the trial was registered in the Chinese Clinical Trials Registry (MR-11-24-048384). All procedures were conducted in accordance with the ethical standards outlined in the Declaration of Helsinki. All participants provided written informed consent.

## Authors contribution

Yi Ye, Kaitong Ma, Xujun Duan and Li Zhang designed the study and developed the framework. Erni Ji contributed to data curation. Yi Ye and Kaitong Ma analyzed the data and wrote the original draft. Maolin Hu, Xiaofen Zong, Zhen Liang, Huang Gan, Bharat B. Biswal, Xujun Duan and Li Zhang contributed to results discussion. Bharat B. Biswal, Xujun Duan and Li Zhang review and editing the manuscript. Zhen Liang, Gan Huang and Li Zhang contributed to funding acquisition. All authors contributed to the final version of the manuscript.

